# It’s all about location: reliance on spatial rather than visual context when trying to remember

**DOI:** 10.1101/2023.02.20.529294

**Authors:** Keren Taub, Shlomit Yuval-Greenberg

## Abstract

When people try to remember visual information, they often move their eyes similarly to encoding. The mechanism underlying this behavior has not yet been fully understood. Specifically, it is unclear whether the purpose of this behavior is to recreate the visual input produced during encoding, or the motor and spatial elements of encoding. In this experiment, participants (N=40) encoded pairs of greyscale objects, overlaying colored squares. During test, participants were asked about objects’ orientation, while presented with squares of the same colors, either at the same location (controlled trials) or switched in their locations (test trials) relative to encoding. Results show that during test trials, participants tended to gaze at the square appearing at the location where the remembered object was previously presented, rather than on the square of the same color. This indicates a superiority of motor and spatial elements of eye movements rather than near-peripheral visual cues.

## Introduction

Our eyes not only allow us to see, but are also involved in various other cognitive tasks. Research has found that people’s eyes often move around when they are attempting to solve mathematical problem, think creatively, make decisions and remember previously learned information (Abeles & Yuval-Greenberg, 2017; Brandt & Stark, 1997; Ferreira et al., 2008; Renkewitz & Jahn, 2010; Spivey & Geng, 2001). Even when the visual information is not relevant to the task, these eye movements still occur.

Eye movements performed during memory retrieval are not arbitrary; rather, findings show that observers tend to reenact the same eye movements they have performed while encoding the information (Brockmole & Irwin, 2005; Johansson & Johansson, 2014; Melcher & Kowler, 2001). This behavior is evident regardless of the type of retrieved information, and has been found for visual objects (Spivey & Geng, 2001b), scenes (Altmann, 2004) and even for facts that were presented auditorily (Scholz et al., 2016). Moreover, it has been found that similarity in eye movements between encoding and retrieval is associated with higher memory performance suggesting that eye movements serve a functional role as retrieval cues (Johansson & Johansson, 2014). However, the mechanism by which gaze aids memory retrieval remains hitherto unknown.

In a previous study, we showed that the benefit of eye movements to memory retrieval is dependent on the visual input (Taub & Yuval-Greenberg, 2023). We found that when most of the visual scenery presented during encoding was blocked from view during retrieval this eliminates the benefit of eye movements to memory. To do this, we placed a large physical occluder that blocked all the near- and most of the far-peripheral input from the participant’s field of view. Following that study, however, it remained unclear which type of visual input contributed to this effect, and specifically whether it is the near peripheral input or the far peripheral input combined with spatial and motor information. As an example, when trying to recall the title of a book that used to be on a specific shelf location, shifting one’s gaze towards that shelf location may improve recall. One hypothesis is that recall is improved because shifting the eyes activates a representation of the spatial index of the book’s location (spatial hypothesis). This representation could be activated by the specific motor action involved in the gaze shift, or by the visual input originating in the far periphery, including the bookshelves and the rest of the room’s layout. Alternatively, a second hypothesis suggests that shifting the gaze enhances memory because it brings to view the items surrounding the recalled book, such as the other books on its shelf, thereby recreating the specific near-peripheral retinal image that was present when that book was previously seen (visual hypothesis).

In this study we examine the contribution of near visual context and spatial location to the effect of eye movements on memory recall. Our goal is to understand why people move their eyes to similar locations during retrieval as they did during encoding: do they try to recreate a similar visual context to the one seen on their fovea during encoding, or do they try and recreate the same spatial location by shifting their eyes to the same position? In the experiment, 40 participants encoded a set of objects presented with different colored squares in the background and were then asked to recall their orientation while presented with the same colored squares, now without objects. During recall, the squares were positioned either in the same positions as during encoding (in “control” trials) or in switched positions (in “test” trials). In control trials, we predicted that participants would look at the location that encapsuled both visual and spatial cues (the same color of the background square and the same spatial location); but in test trials, participants could gain from looking either at the original location of the object, recreating the same spatial cue, or at the congruent color, recreating the same near-peripheral visual context that accompanied the target object during encoding. Findings of this experiment would disentangle the contribution of the different components of eye movements, and explain the mechanism underlying the use of eye movements during memory retrieval.

## Methods

### Participants

A total of 40 individuals participated in the experiment (27 females, age range 19-29, Mean age = 24.08, SD age = 1.85). Sample size was chosen based on a power analysis using Gpower 3.1 (Faul et al., 2007), with an estimation of a medium effect size (Cohen’s d=0.5), with significance level of 0.05 and power of 0.8. Power estimation indicated a required sample size of at least N=34. Participants were students of Tel Aviv University, that were recruited through an online subjects-recruitment system, and received course credit for their participation. All participants had normal or corrected-to-normal vision and were native Hebrew speakers. Participants signed an informed consent prior to the experiment. The experiment was approved by the ethical committee of Tel Aviv University and the School of Psychological Sciences.

### Stimuli

Stimuli consisted of 128 grayscale drawings of objects, taken from Snodgrass and Vanderwart’s object pictorial set (Snodgrass & Vanderwart, 1980) and Multipic database (Duñabeitia et al., 2018; http://www.bcbl.eu/databases/multipic). Three quarters of these drawings were included in three blocks of the main experiment, and one quarter was included in an extra block that was intended to test participants memory for colors. Stimuli for this experiment were selected based on preliminary research in which 200 objects were presented to 20 native Hebrew participants who were asked to indicate the name and orientation of each object. Only objects on whose identity and orientation there was a wide consensus of at least 90% of participants, were included in the main experiment. High acquittance with the objects’ names and correct identification of their orientation were also validated for each participant of the main experiment at the end of the session. Objects whose names or orientations were not correctly recognized by a specific participant were excluded from analysis for that participant.

The drawings of the objects were presented on top of colored-squares, of 7 visual degrees horizontally and 9 visual degrees vertically. The colors of the squares were taken from a group of six same-luminance colors that were notably distinguished from one another (in RGB: red: 255, 0, 0; yellow: 255, 255, 0; green: 0, 255, 0; light blue: 0, 255, 255; pink/purple: 255, 0, 255; blue: 0, 0, 255).

### Procedure

Prior to the main experiment, participants completed a short training session with six objects (three pairs) in the presence of the experimenter, to ensure that they understood the task. Thet were told that the main experiment included four blocks, each starting with an encoding phase (100 seconds), followed by a test phase (lasting for 5-6 minutes, depending on participants’ speed of response) in which their memory regarding the objects will be tested. In practice, only the first three blocks included an encoding and test phases, and the fourth included a different task, aiming to test their memory for colors.

The encoding phase of each of the three main-experiment blocks consisted of 16 trials, each presenting two objects, with a total of 32 objects per block. Each encoding trial started with 1 second of a gray screen followed by the presentation of two colored squares (sizes 7×9 visual degrees), presented 4 visual degrees to the left and the right of the screen’s center. Following 100 ms, two object drawings appeared on top of the squares and at their centers (see **Figure 1**). After another 5 seconds, the objects disappeared and the squares remained for another 100 ms before they disappeared as well and the next trial began. In the encoding phase of each block, half of the objects (randomly picked) were oriented to the right and half to the left. Participants were asked to memorize objects’ orientation and received no instructions regarding the colored squares. At the end of the encoding phase appeared a message on the screen notified participants of the upcoming test, that started after a pause of 15 seconds.

**Figure 1:**
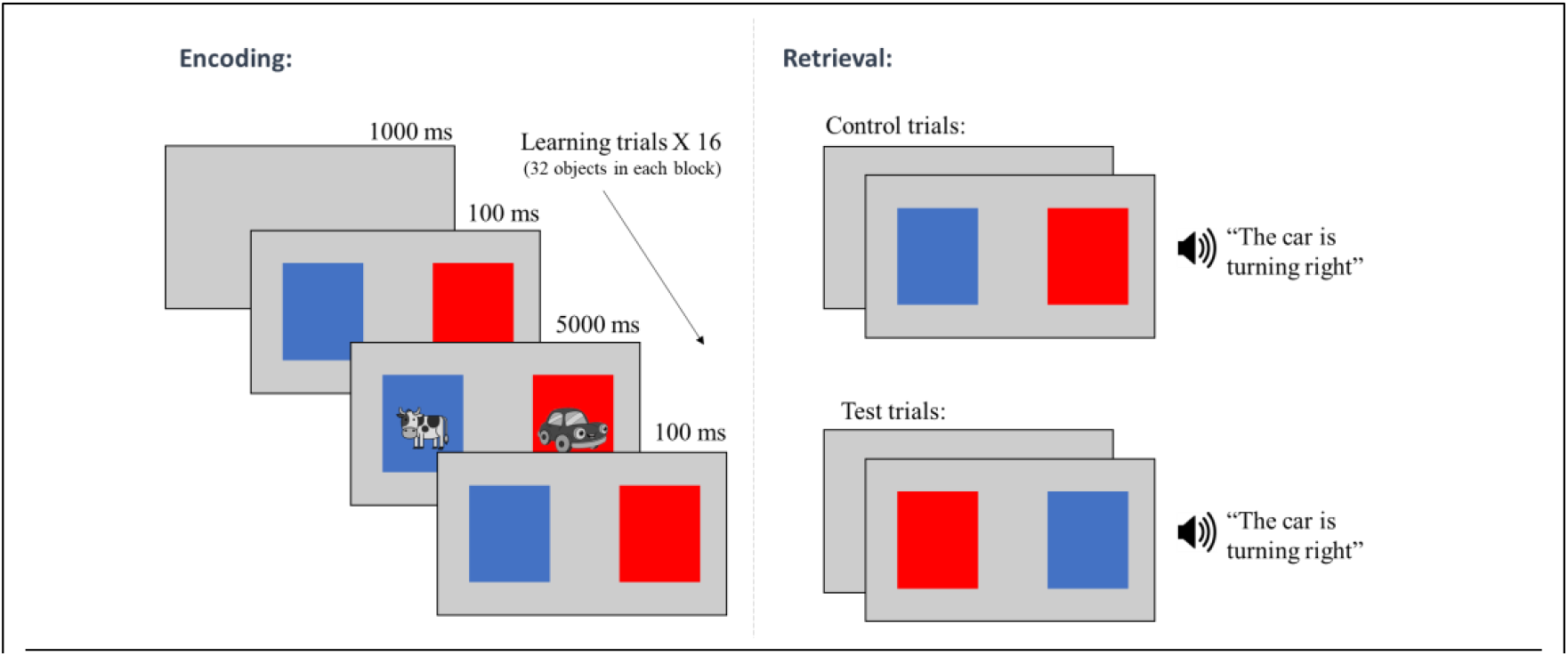
An encoding trial with its to possibilities during test. During encoding, participants first saw two colored squares, then two greyscale objects appeared and participants were requested to memorize their orientation. During test, participants heard statements regarding objects orientation, while seeing the same two colored-squares: either in the same locations (control trials) or in opposite locations (test trials).

In the test phase of each of the three main-experiment blocks, participants were presented with two colored squares at the same locations as in the encoding phase. Each trial corresponded to a trial of the encoding phase and included squares of the same colors as in that trial. The colors appeared either in the same side as in the original trial (‘control trial’) or in the opposite side (‘test trial’). Participants listened to pre-recorded statements describing an object’s orientation (e.g. “The zebra is turning right”) and were instructed to respond by pressing one of six keys, indicating both their response to the statement (True / False) and their confidence level (on a scale of 1-3). When participants pressed a key, the trial ended and the next trial began. At the end of each block, participants received feedback on their performance including their percent of correct responses.

The last block of the experimental session included a different task. The initial encoding phase was as in the main experimental blocks, but before test, new instructions appeared on screen, and the experimenter walked in to ensure that the participants understood the new instructions. In this test phase participants were presented, at each trial, with one of the objects that were previously viewed, placed in the middle of the screen. The background of this object was a square colored in one color on the top half and another color on the bottom half. The colors were the same colors that were presented with that object when it first appeared in the encoding phase of the main experiment blocks. One color was the color that consisted the background of the target object in its initial presentation, and the other color was the one that consisted the background of its pair object during the initial presentation. Participants were requested to recall which color appeared with the target object when it was initially presented and to report that color using the up and down arrows of the keyboard. In half of the trials, the correct color appeared at the top part, and in the other half it appeared on the bottom part.

### Apparatus and Eye tracking

Participants were seated in a sound-attenuated room at a distance of 100 cm from a 24 inch ASUS VG248QE LCD screen, with their head rested on a chin-rest. Binocular eye-movements were monitored binocularly using a remote infrared video-oculographic system (EyeLink 1000 Plus, SR Research Ltd, Ontario, Canada), with a sampling-rate of 1000 Hz, spatial resolution <0.01° and average accuracy of 0.25° - 0.5° when using a head-rest, as reported by the manufacturer. At the beginning of each block, participants completed a 9-point eye tracking calibration procedure performed with the same illumination conditions as the main experiment. Calibration was accepted when validation confirmed that fixation errors were of no more than 0.2°.

### Analysis

Eye movements analysis was performed using Matlab R2021b. Fixations were included in the dwell time analysis if they were longer than 100 ms and landed within the borders of the screen. Trials with no fixations within screen limits were omitted from analysis, and so were trials presenting objects that participants wrongly identified their name or orientation (percent of trials that were included in the analysis: mean 93.7%, SD: 5.05%, range across participants 70.83%-96.88%). In each trial, a fixation was defined as congruent with the original location (“congruent fixation”) if it was performed during a test phase trial and landed within the side of the screen where the target object was originally presented during encoding. Fixations that landed on the other side of the screen, were referred to as “incongruent fixations”.

## Results

### Accuracy rates

The average accuracy rate across conditions was 74.68% (SD=10.76%, range: 55.21%-96.88%), with no significance difference in performance between test and control trials (t(39)=0.518, p=0.608). Results for accuracy according to condition are presented in **Table 1**.

**Table 1.** Accuracy results of memory for orientation, divided according to condition.

|  | <u>Test trials</u> | <u>Control trials</u> |
| --- | --- | --- |
| Mean<br>(SD) | 74.12%<br>(13.22%) | 74.92%<br>(10.43%) |
| Range | 45.83%-97.92% | 52.94%-100% |

### Gaze behavior

We conducted a 2×2 ANOVA with independent variables Condition (test trials / control) and Spatial-congruity (congruent location / incongruent location) with gaze dwell time as the independent measurement. We found main effect for screen side, with participants looking more at the congruent side, meaning looking towards the same location as in encoding, in both test and control trials (F(1,39)=29.88, p<0.001). Simple effects analysis revealed this behavior was significant in both control trials and test trials (control trials: t(77.88)=4.112, p<0.001; test trials: t(77.88)=3.769, p<0.001, see **Figure 2**). Note that, in the test trials, the location that was spatially-congruent was always incongruent in its color (the background color of the original target is always the color of the square at the incongruent location). Therefore, finding a simple effect of congruity in the test condition indicates that gaze tended to dwell over the congruent side rather than on the congruent color. We found no main effect for Test condition (F(1,39)<1, p=∼1), and no interaction between Screen side and Test condition (F(1,39)=0.061, p=0.81).

**Figure 2:**
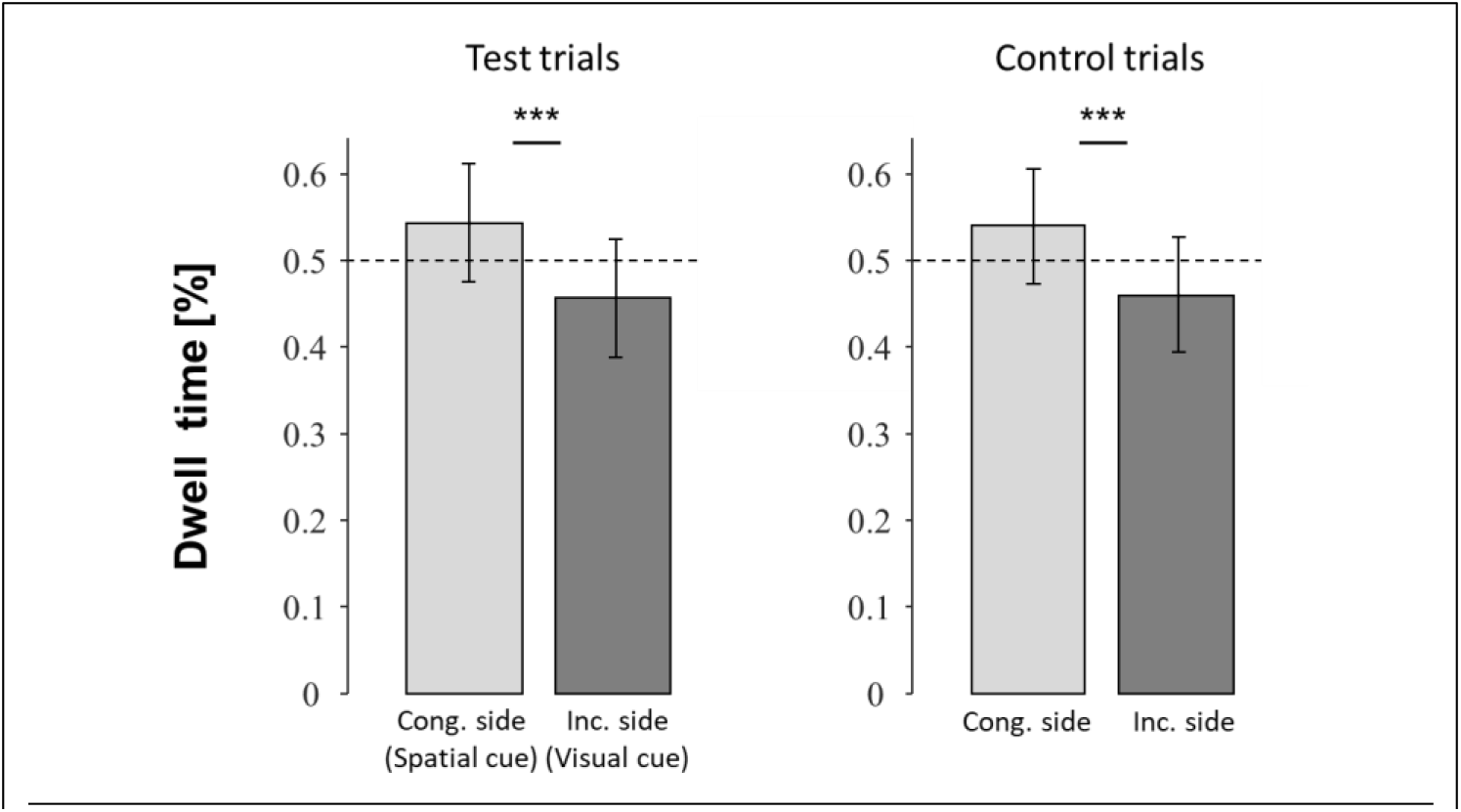
Dwell time analysis revealed that for both test trials and control trials, participants preferred looking towards the part of the screen where the target object originally appeared. *** p<0.001

### Retrieval performance according to gaze congruency

To test for the effect of gaze location on accuracy, trials were divided to spatially-congruent trials (trials in which more than 50% of fixation time was on the side of object original location) and spatially-incongruent trials (<50%). We performed a 2×2 ANOVA to test accuracy according to Gaze spatial-congruency (congruent / incongruent) and Condition (test / control).

We found no evidence for a main effects (Condition: F(1,39)=0.11, p=0.92; Gaze spatial-congruency: F(1,39)=0.34, p=0.56) or an interaction (F(1,39)=1.17, p=0.29).

Follow-up Bayesian tests confirmed moderate support for the null hypothesis for both main effects (main effect of condition: BF_01_=4.808, error=1.358%; main effect of gaze congruency: BF_01_=4.535, error=2.533%) and strong support for the null hypothesis of the interaction (BF_01_=43.374, error=1.885%). These findings indicate that gazing at either the spatially-congruent and the color-congruent location, did not enhance memory performance. These results are presented in **Figure 3**.

**Figure 3:**
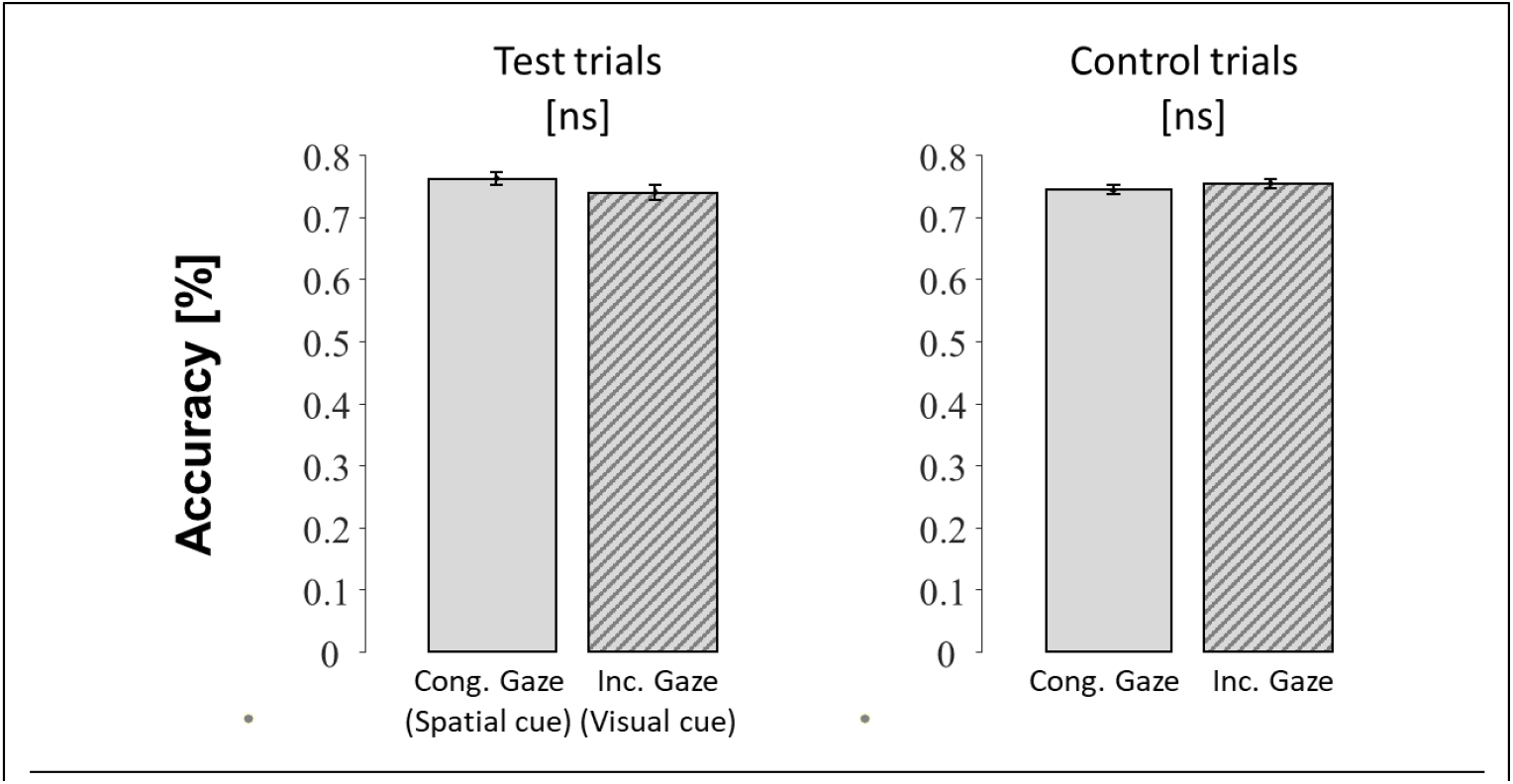
Accuracy analysis revealed that neither Test condition, nor Gaze congruency, affected performance in the test.

### Memory for colors

Gazing at a square of the same color as the target’s background did not enhance the recall of that target. It could be suggested that since color in this design was task-irrelevant it was ignored and not encoded. To examine this possibility, we performed the last block in which we assessed the participants’ memory for the combination of objects and square colors. In this task participants were asked to, explicitly, report the color of the background presented with each object that they viewed. We conducted a one sample t-test and found that accuracy rates were higher than chance level performance of 50% (t(39)=2.171, p=0.036). Single-subjects results for this analysis are depicted in **Figure 4**.

**Figure 4:**
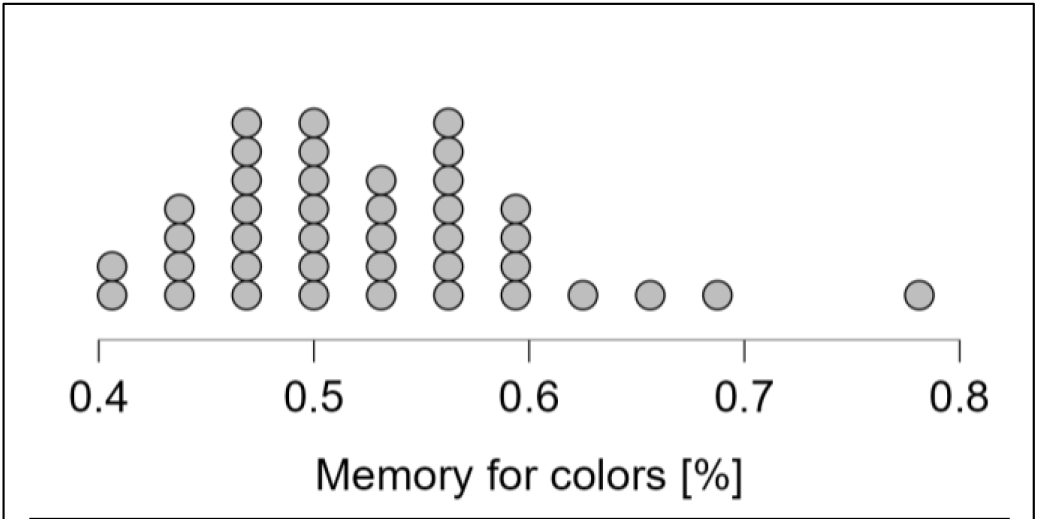
Distribution of participants results in the memory test for colors. Chance level results are 50%, indicating the most of the participants succeeded more than chance level.

## Discussion

Consistently with previous studies we found that people tend to look during retrieval at the same locations where they fixated during encoding. Our results demonstrate that this behavior is robust even when the near-peripheral visual input during retrieval does not match the one viewed during encoding. Nevertheless, the tendency to gaze at the original location of an encoded object rather than at the location that is more visually consistent with it, was found to be unbeneficial to memory performance, thus raising questions regarding its functionality.

The finding that eye movements performed during retrieval are largely similar to those performed during encoding, even though visual information in the near-periphery is no longer available, is consistent with previous studies (Brockmole & Irwin, 2005; Johansson & Johansson, 2014; Melcher & Kowler, 2001). However, in contrast to previous studies, here we dissociated the near-peripheral visual input that accompanied the encoding of an object from the object’s location, and consequently from the far-peripheral visual input. This revealed that participants preferred gazing towards the original location rather than gazing towards the colored square that surrounded the target object during encoding.

While this behavior could indicate a general tendency in retrieval towards the spatial resemblance over the resemblance in the visual input, it is worth noting that this experiment only made use of the visual input the was in close proximity to the fovea, and only manipulated the color in this near-periphery area. Moreover, it should be taken into consideration that while a meaningful part of the visual image on the retina was the color of the square in the background of each object, there were other visual details in the periphery that affected visual context, such as the edges of the screen and other visual elements in the room, that remained in place during test and therefore were not part of the shift we tried to create in the visual context. That is, while the squares were oppositely colored during test trials, other visual elements in the visual field remain the same, keeping the original location of encoding not only the source of spatial reconstruction, but also as a location that withhold at least some visual information from encoding. To switch the locations of full visual context there is need in producing a unique setting, possibly using VR, to eliminate all other visual cues and ensure that the location is the only element that changes. It is likely that greater similarity in visual input and over a larger area of the visual field would have made the visual context more useful during retrieval.

It is further possible that a different visual context than colors would have alter the effect on memory performance, as the visual resemblance in this experiment relied solely on the color on the background square. Other visual elements, even if located in the near periphery, may have had a greater impact on encoding and therefore would have been more meaningful if revisited during retrieval. For example, if near-periphery visual input was more relevant, more complex or more salient – factors that were found to affect the encoding of memories – memory performance might have altered (Chun & Turk-Browne, 2007; Constant & Liesefeld, n.d.; Kemps, 2001).

Another alternative explanation for these findings is that the specific memory task that was used may have encouraged the reliance on spatial rather than visual information. In this task participants were asked to encode objects’ orientation: left or right, a task with a spatial component. It is possible that this task enhanced the encoding of spatial rather than the visual information. It could be the case that a different task, for example one that focuses on color, would have resulted in a different outcome.

Some previous studies, including our own (Taub et al., 2022; Taub & Yuval-Greenberg, 2023), have shown that the tendency to look at similar locations during encoding and retrieval is beneficial for memory performance (Bochynska & Laeng, 2015; Johansson & Johansson, 2014; Spivey, 2008). However, in the present study, we have not found an enhancement effect. Even in control trials, in which the color-congruent location matched the spatially-congruent location, there was no improvement in recall performance in trial in which the gaze dwelled more on the congruent side.

There are a few possible hypotheses as to why in the current settings memory enhancement did not occur following a congruent gaze. Most likely, this lack of effect is related to the fact that during encoding, in each block, participants were asked to memorize 32 different objects, while all objects appeared in either one of two locations. This means, that locations were used repeatedly by different objects and were not highly linked to specific object by the end of the encoding. As colors were also not object-specific, both visual and spatial cues were not unique. This could be explained in light of the “cue overload” theory, that suggested that a retrieval cue may become less effective when it is shared by multiple items (Watkins & Watkins, 1975). Due to this concept, the repetition among colors and locations impaired their effectiveness as possible cues in the current research. Use of distinct locations and unique visual resemblance might have turn gaze congruency to be more efficient in memory retrieval.

## Conclusion

The current research serves as evidence for the human tendency to recreate similar gaze positions between encoding and retrieval when trying to retrieve previously learned information. Our results demonstrate the strength of this behavior even when it is not accompanied by a similar visual input that could be found in a different position, and emphasize that this behavior take place even when it is unbeneficial to memory performance. This implies that the ability to gain from retrieval cues related to eye movements does not occur in all conditions, and suggests that future research in the field could identify what are the terms that enable people to gain from these cues.

## Acknowledgements

This research was funded by the Israel Science Foundation, grant 1960/19 to S.Y-G and by the Ariane de Rothschild Women Doctoral scholarship to K.T.

